## Supplementary material for "Lack of tetrodotoxin analogues and individual metabolomic profiling of the cryptic frog *Colostethus imbricolus*": S1 File

Figure S1. Graphical abstract from this study. Created in [https://BioRender.com](https://biorender.com)

Figure S2. Base peak chromatograms (BPC) (to the left) and the corresponding MS/MS fragmentation patterns (to the right) from target masses of interest in the preliminary analysis of *C. imbricolus*. Two instrumental replicates of the same sample are presented for each ion (r001 and r002). The most intense molecular feature (178.1342) and eight target masses of TTX analogues (3 pages). *The 20-minute gradient on the SB-CN column was employed in these analyses.

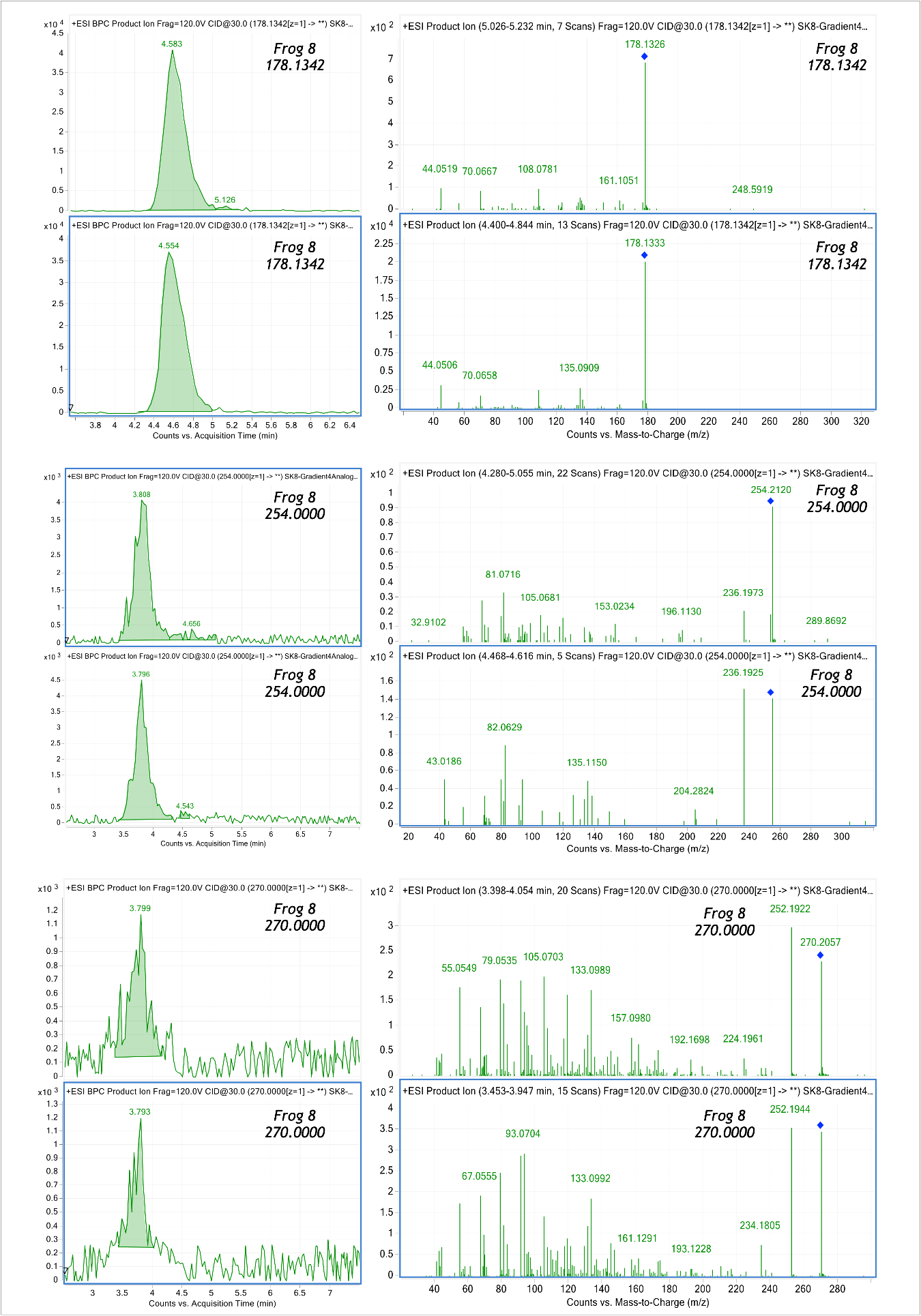

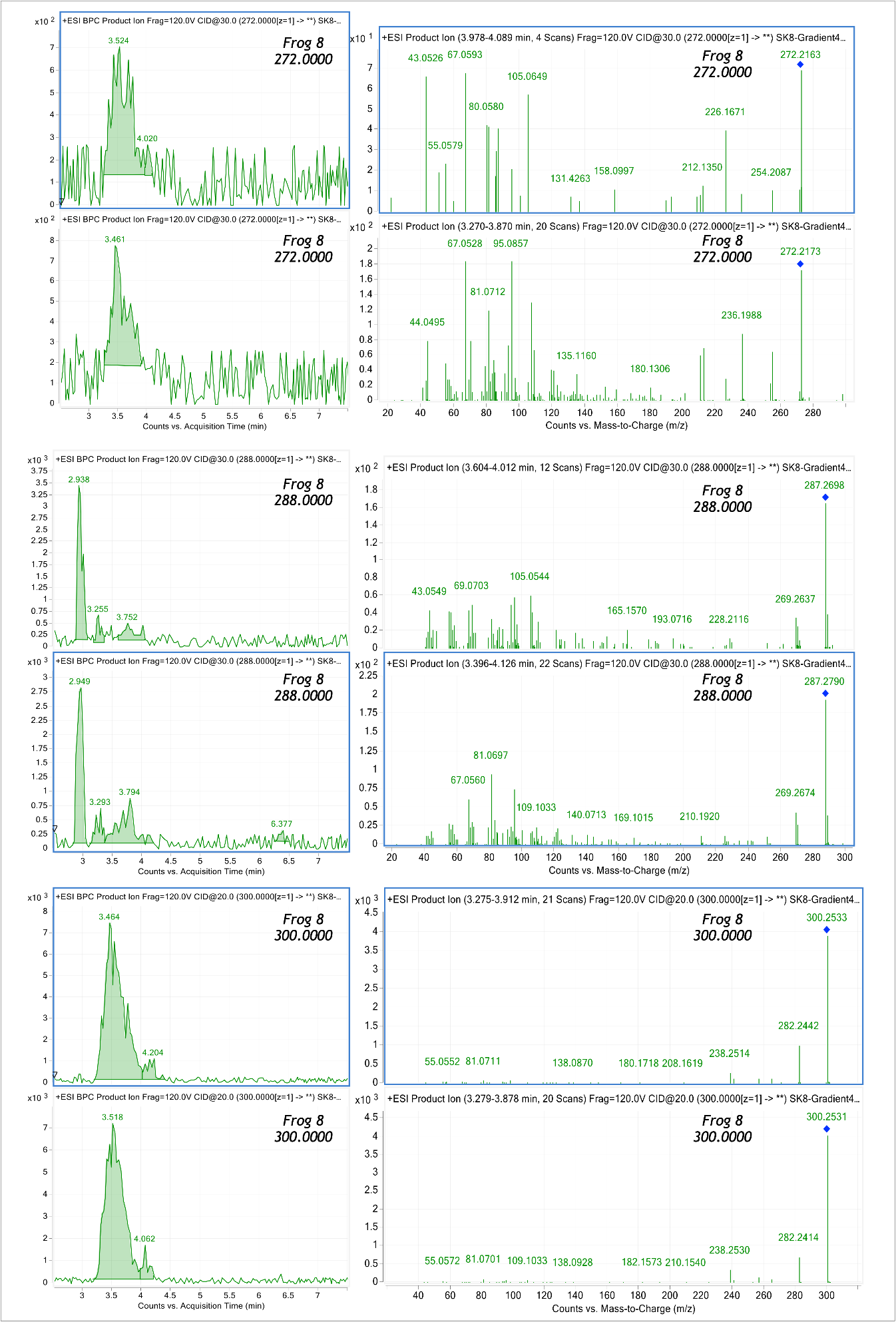

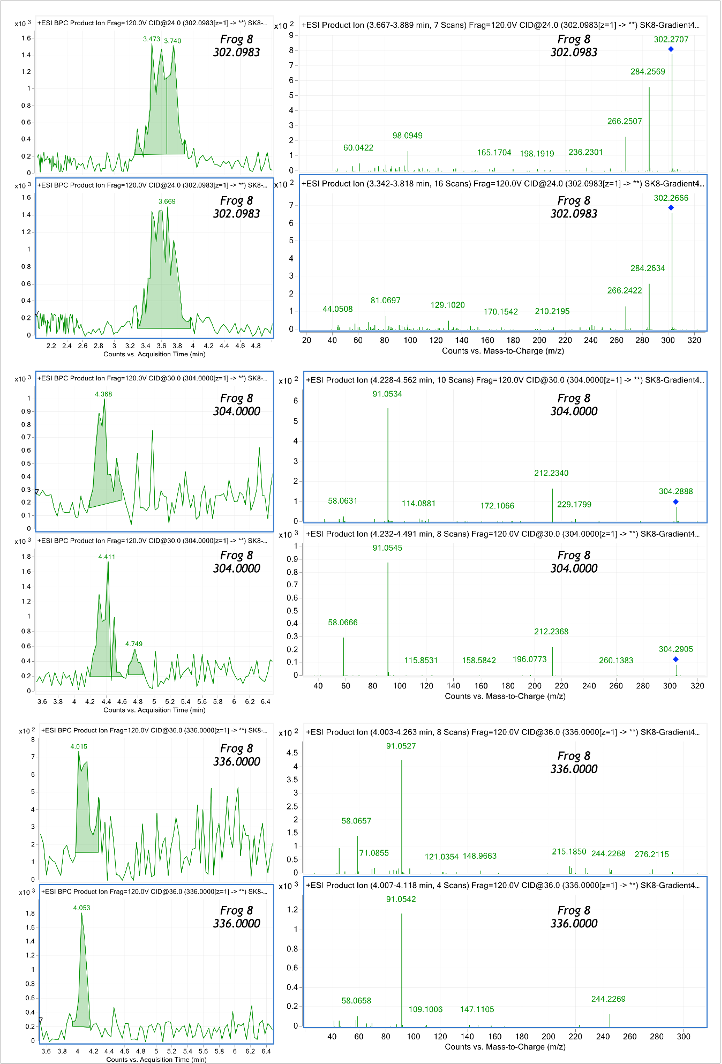

Figure S3. Chromatographic separation of TTX employing a chromatographic method of 20 minutes in SB-CN column. A. Extracted ion chromatogram (EIC) from accurate mass of TTX (320.1088) in a blank of MeOH and three instrumental replicates of 10 ppm TTX solutions. B. Verification of deconvolution process of TTX molecular formula prediction. C. MS/MS fragmentation pattern of TTX. r001, r002 and r003 correspond to instrumental replicates of the same sample.

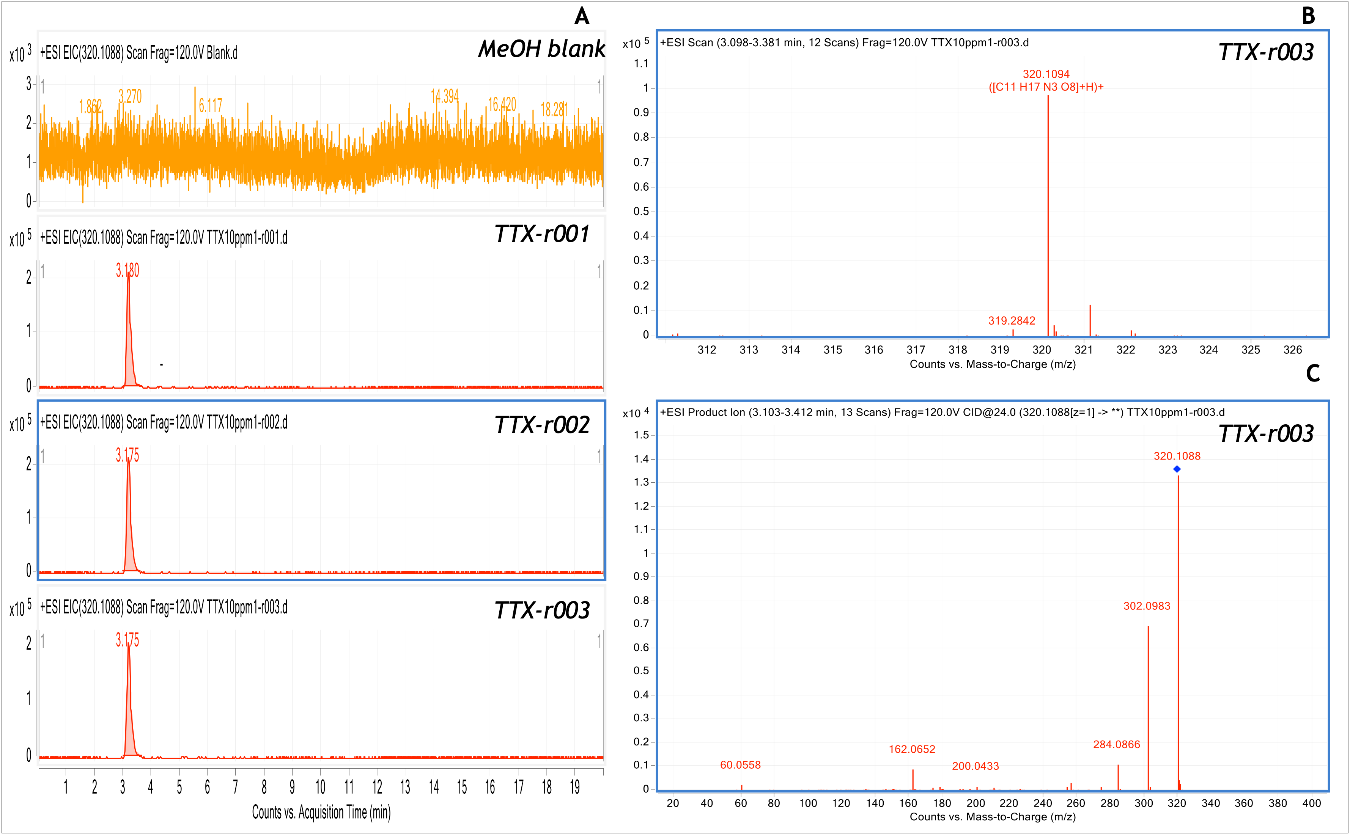
